# Quaternionic Assessment of EEG Traces on Nervous Multidimensional Hyperspheres

**DOI:** 10.1101/2020.03.05.979062

**Authors:** Arturo Tozzi, James F. Peters, Norbert Jausovec, Irina Legchenkova, Edward Bormashenko

## Abstract

The nervous activity of the brain takes place in higher-dimensional functional spaces. Indeed, recent claims advocate that the brain might be equipped with a phase space displaying four spatial dimensions plus time, instead of the classical three plus time. This suggests the possibility to investigate global visualization methods for exploiting four-dimensional maps of real experimental data sets. Here we asked whether, starting from the conventional neuro-data available in three dimensions plus time, it is feasible to find an operational procedure to describe the corresponding four-dimensional trajectories. In particular, we used quaternion orthographic projections for the assessment of electroencephalographic traces (EEG) from scalp locations. This approach makes it possible to map three-dimensional EEG traces to the surface of a four-dimensional hypersphere, which has an important advantage, since quaternionic networks make it feasible to enlighten temporally far apart nervous trajectories equipped with the same features, such as the same frequency or amplitude of electric oscillations. This leads to an incisive operational assessment of symmetries, dualities and matching descriptions hidden in the very structure of complex neuro-data signals.

---

The nervous activity of the central nervous system takes place in dimensions higher than the conventional spatial three plus time. This counterintuitive claim, at first proposed by Tozzi and Peters (2016), has been put forward in terms of simplicial complexes encompassing synaptic connections (Reimann et al., 2017), of neural codes for navigating cognition (Bellmund et al., 2018), of rhythm and synchrony in cortical network models (Chariker et al., 2018). It has been stated that “invariances or symmetries afforded by projections onto high dimensional spaces… may not reveal themselves through local scrutiny of the surface data acquired from the brain in action, but may require” the use of higher-dimensional manifolds (Friston, 2017). For a reviewon the multidimensional brain, see Tozzi (2019). In its early formulation, the higher-dimensional nervous trajectories have been described as occurring in the four spatial dimensions of a genus-zero hypersphere S^3^, or of a genus-one Clifford torus (Tozzi and Peters, 2016; Tozzi et al., 2017). These authors suggest that nervous activities are embedded in an imperceptible fourth spatial dimension, in particular brain functions such as mindwandering and memory retrieval (Peters et al., 2017).

Here we ask: starting from the conventional neuro-data available in three dimensions plus time, does there exist an operational procedure to describe the corresponding four-dimensional trajectories? Is it feasible to assess in higher dimensions the three-dimensional paths detected during standard experimental procedures? Here we describe a viable option: the projection of three-dimensional data achieved from real series to the peculiar manifold of a four-dimensional hypersphere. In particular, we aim to map electroencephalographic (EEG) electric oscillations to an S^3^ hypersphere, or in other words, to achieve orthographic projections of brain signal patches via quaternions.

The development of quaternions (Tate, 1867), first proposed by Hamilton (1843), has been quite slow, due to the painstaking issue of the non-commutativity of quaternion multiplicity. Nevertheless, in recent years, novel improvements to avoid such non-commutativity gave to quaternions the possibility to achieve useful results in attitude control, quantum mechanics, computer graphics, and also in neural network research and modeling patterns of EEG signals. Quaternion-based signal analysis techniques were used to extract features related to motor imagery in various mental states, allowing the classificationof signals in decision trees, support vector machine and k-nearest neighbour techniques (Batres-Mendoza et al., 2016). Li and Wang (2018) assessed drive-response synchronization for quaternion-valued shunting inhibitory cellular neural networks with mixed delays. The occurrence of almost periodic solutions, achieved through the Banach fixed point theorem, led to novel state-feedback controllers to ensure global exponential synchronization. The last, but not the least, Enshaeifar et al. (2016) introduced a novel quaternion-valued singular spectrum analysis for multichannel analysis of EEG, aiming to provide an effort to solve the excruciating problem of noise in the extracted sources.

After exploring the possibility to use global visualization methods for exploiting quaternion maps of EEG traces, we will show in the sequel how a quaternionic description of brain activity, apart from the assessment of the multidimensional brain, provides other operational benefits that greatly improve the extraction of useful information and the evaluation of neuro-data.

## MATERIALS AND METHODS

We aimed to map two- and three-dimensional EEG oscillations extracted from real neurodata series to a four-dimensional S^3^ hypersphere. To achieve our goal, we used four-dimensional quaternions to represent orthographic projections of brain signals’ images onto the more manageable surface of a three-dimensional sphere.

### EEG traces

We analyzed EEG traces of subjects at rest. EEG was recorded using a Quick-Cap with sintered (Silver/Silver Chloride; 8mm diameter) electrodes. Using the Ten-twenty Electrode Placement System of the International Federation, the EEG activity was monitored over nineteen scalp locations (Fp1, Fp2, F3, F4, F7, F8, T3,T4, T5, T6, C3, C4, P3, P4, O1, O2, Fz, Cz and Pz). ll leads were referenced to linked mastoids (A1 and A2), and a ground electrode was applied to the forehead. Additionally, vertical eye movements were recorded with electrodes placed above and below the left eye. The digital EEG data acquisition and analysis system (SynAmps) had a bandpass of 0.15-100.0 Hz. At cutoff frequencies, the voltage gain was approximately –6dB. The 19 EEG traces were digitized online at 1000 Hz with a gain of 1000 (resolution of 084μV/bit in a 16 bit A to D conversion), and stored on a hard disk. Epochs were automatically screened for artifacts. All epochs showing amplitudes above +/-50 microV (less than 3%) were excluded, to avoid that the traces could be artifacts of the visualizing algorithm while plotting the EEG power maps. The EEG study was done according with Declaration of Helsinki and was approved by the Ethics Committee of the University of Maribor, Slovenia.

The traces assessed in this study were extracted from three brain areas, corresponding to the central electrodes Cz, Fz, Pz. These peculiar locations were chosen because of the lower occurrence of oscillatory artifacts in these three scalp areas.

### Fourier analysis

A Fourier analysis was performed to define the underlying frequencies of the EEG data sets. To perform this analysis, a corresponding MATLAB based software was designed. The frequencies were obtained with resolution 0.25 Hz. It is well known that the brain activity described by Fourier analysis can be tackled in terms of scale-free dynamics (Pritchard, 1992; Fingelkurts et al., 2010; Buszaki and Watson, 2012). Indeed, cortical electric oscillations observed at many spatiotemporal scales exhibit a frequency spectrum that displays a scale-invariant behavior, characterized by a power spectrum, a frequency and an exponent that equals the negative slope of the line in a log power versus frequency plot (Van de Ville et al., 2010; Jirsa et al., 2014; De Arcangelis and Herrmann, 2010).

### Quaternion maps

Anassortment of possible quaternion-driven variants and higher-dimensional visual representation strategies is available, resulting in very complex quaternion maps. The two most widely used classes of quaternion visualization are based on visual geometric context and parallel coordinates (Hanson and Thakur, 2012). In this paper, we use the more manageable geometric view. Because the mathematical procedures have been thoroughly described by many authors, here we favour an approach for non technical readers, based on plainillustrations.The website https://quaternions.online/provides an introduction to 3D quaternion graphics. See also, for an affordable treatment of quaternions, Hart (2014).

In an EEG trace (**Figure 1A**), every point inside the plot displaying amplitude versus time can be described by three vectors lying each one on a different spatial coordinate: x, y or z (say, e.g., green, red and blue arrows). We will term these three vectors “triad”. In the peculiar case of an EEG plot, in which the trace is two-dimensional, we assume that one of the three coordinates equals zero. **Figure 1B** illustrates, for sake of clarity, a triad embedded in a cubic manifold. Quaternion trajectories in three dimensions can be depicted as arcs moving inside a three-dimensional frame encompassing the axes x, y, and z (**Figure 1C**). The movement of the arc is dictated by the constraints of the axis rotation and the Euler angle (Hart, 2014). Every single point of the EEG trace can be projected to a hyperspheric manifold, either using punctiform (**Figure 1D**), or uniform (**Figure 1F**) orthographic projections. **Figure 2** summarizes the quaternionic movements of the triad that are allowed in three-dimensional projections.

**Figure 1.**
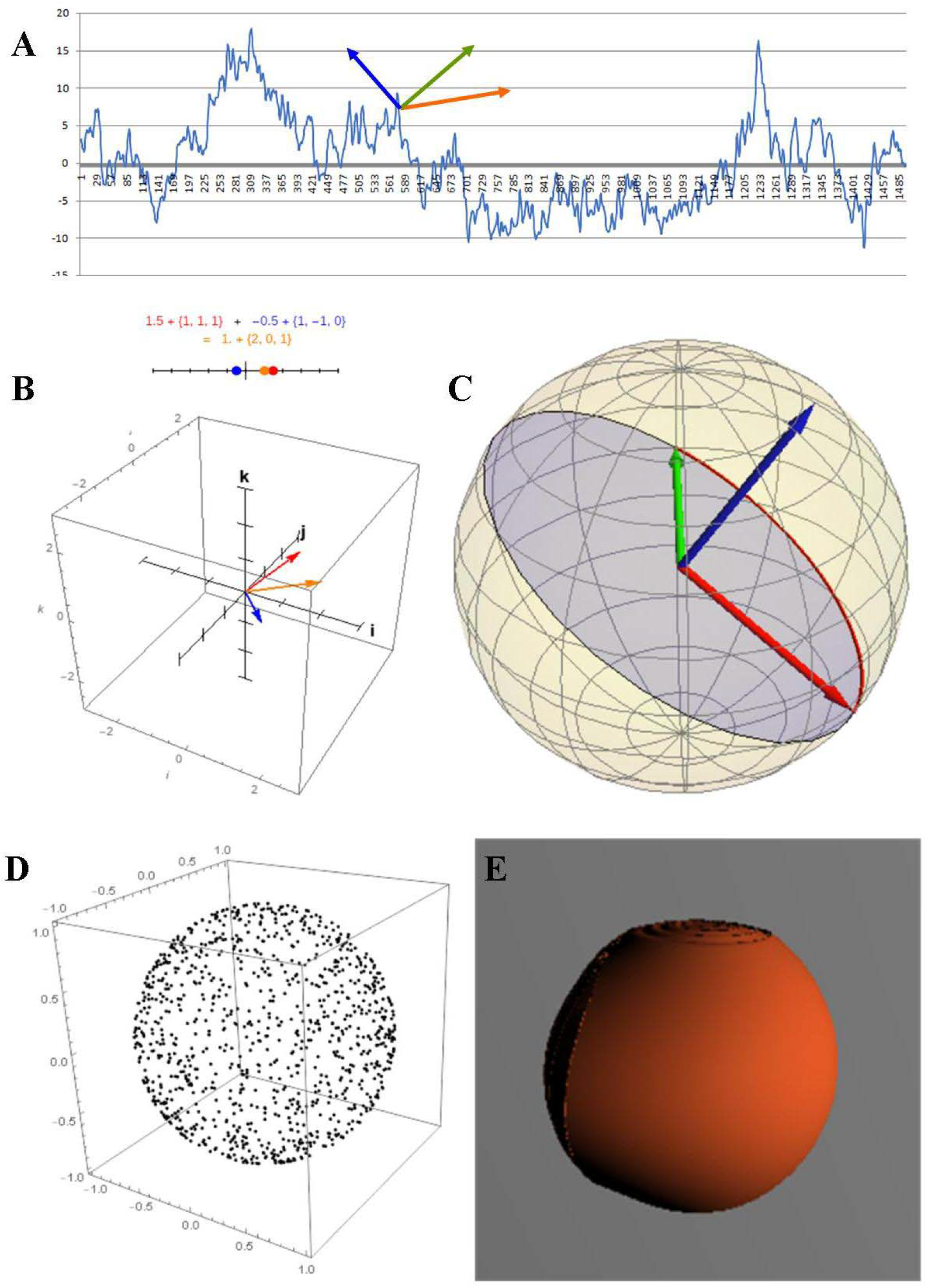
Quaternion mappings from a three-dimensional wave to a four-dimensional hypersphere. **Figure 1A**: in a plot of time in milliseconds (axis x) versus amplitude in mV (axis y), the mean of the oscillations from three brain locations is depicted. The chosen brain areas correspond to the central electrodes Cz, Fz, Pz, were the artifacts are lower. **Figure 1B**: a barebones quaternionic projection relative to the three vectors. **Figure 1C**: three-dimensional quaternion projection inside a sphere representing the brain. **Figure 1D**: punctiform orthographic projection of a brain signal spike (amplitude) onto the surface of a sphere via quaternion sphere. **Figure 1E**: uniform orthographic projection of brain signals onto the surface of a sphere, achieved after projecting brain signal vectors via quaternionic maps.

**Figure 2.**
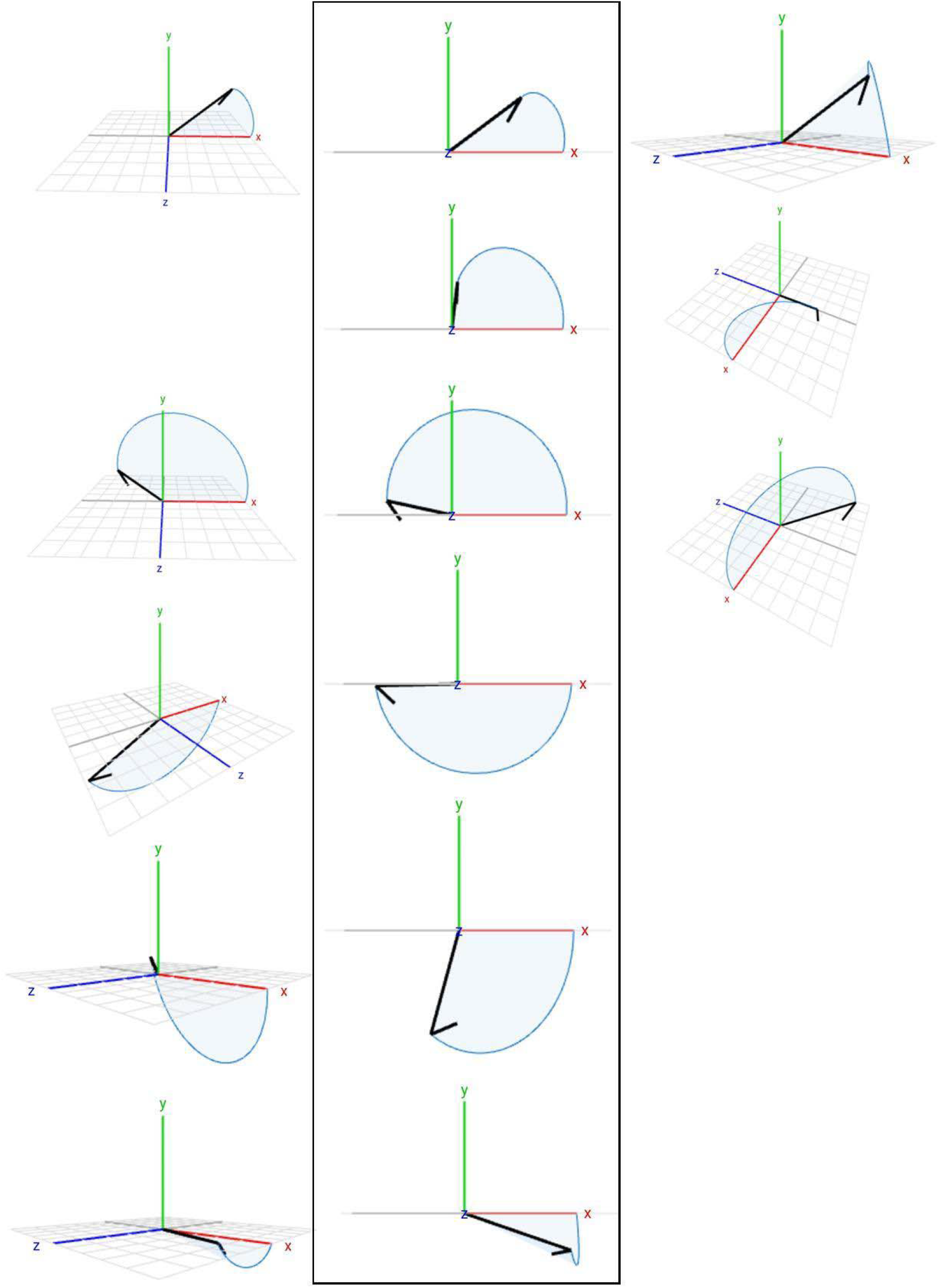
Quaternion trajectories in three dimensions. The central row displays the temporal progression of the trajectories that are visible when the hidden axis z is undetectable by the observer. The pictures on the left and on the right illustrate the same movements described in the central rows, but, for sake of clarity, the axes x, y, and z are placed with different angles to fully appreciate the whole three-dimensional view. Modified from: https://quaternions.online/.

In more technical terms, a map of a glome equipped with Sp(1) or SU(2) Lie groups can be projected onto a threedimensional surface (Tozzi and Peters, 2016). The 3-sphere is parallelizable as a differentiable manifold, with a principal U(1) bundle over the 2-sphere. The 3-sphere’s Lie group structure is Sp(1), which is equipped with quaternionic 1X1 unitary matrices(Odzemir 2013). The symmetries of the quaternionic manifold display a number system similar to the complex numbers, but with three imaginary quantities, instead of just one (Lemaître 1948). We can state that orthographic projections derived from a sphere with center (0,0,0) with unit vectors *α, β, γ*, express the excess in any shape as an orthographic projection on the surface of a sphere (Tait 1867). We express as an arc *DF* and an angle on the surface of the sphere the quaternion *βα*^-1^*γ*, i.e., *DF* = *βα*^-1^*γ*.

Summarizing, the use of quaternions makes it possible to utilize the three spatial coordinates of every point inside a threedimensional plot to become a single point inside a four-dimensional hypersphere. In the sequel, we will describe how brain signal patches derived from Fourier analysis can be represented geometrically as inputs to a quaternion. Indeed, a quaternion provides a cumulative fractal view of multiple brain signals, described in terms of orthogonal projections on a surface of the brain viewed as a sphere.

## RESULTS

We built the two-dimensional plot of an EEG Fourier analysis (frequency in Hz vs. amplitude in mV) (**Figure 3A**), and achieved the corresponding three-dimensional plot (**Figure 3B**). Every brain signal assessable in terms of threedimensional vectors (triads)was projected to four-dimensional quaternionic mappings, depicted as orthographic projections in a three-dimensional space (**Figure 3C**). In other words, such ortographic projections are represented by points on threedimensional arcs located inside a sphere, the latter standing for the phase space of the brain where electric EEG oscillations take place (**Figure 3C**). Note that the triads with matching description in the three-dimensional plot (see the two triads depicted in **Figure 3B**) converge towards the same area of the four-dimensional hypersphere (the yellow arrows and the single triad depicted in **Figure 3C**): this means that temporally distant three-dimensional trajectories equipped with the same features (say, the same amplitude or the same frequency) are easily enlightened in four dimensions, because they come to be very close inside the quaternionic map.

**Figure 3.**
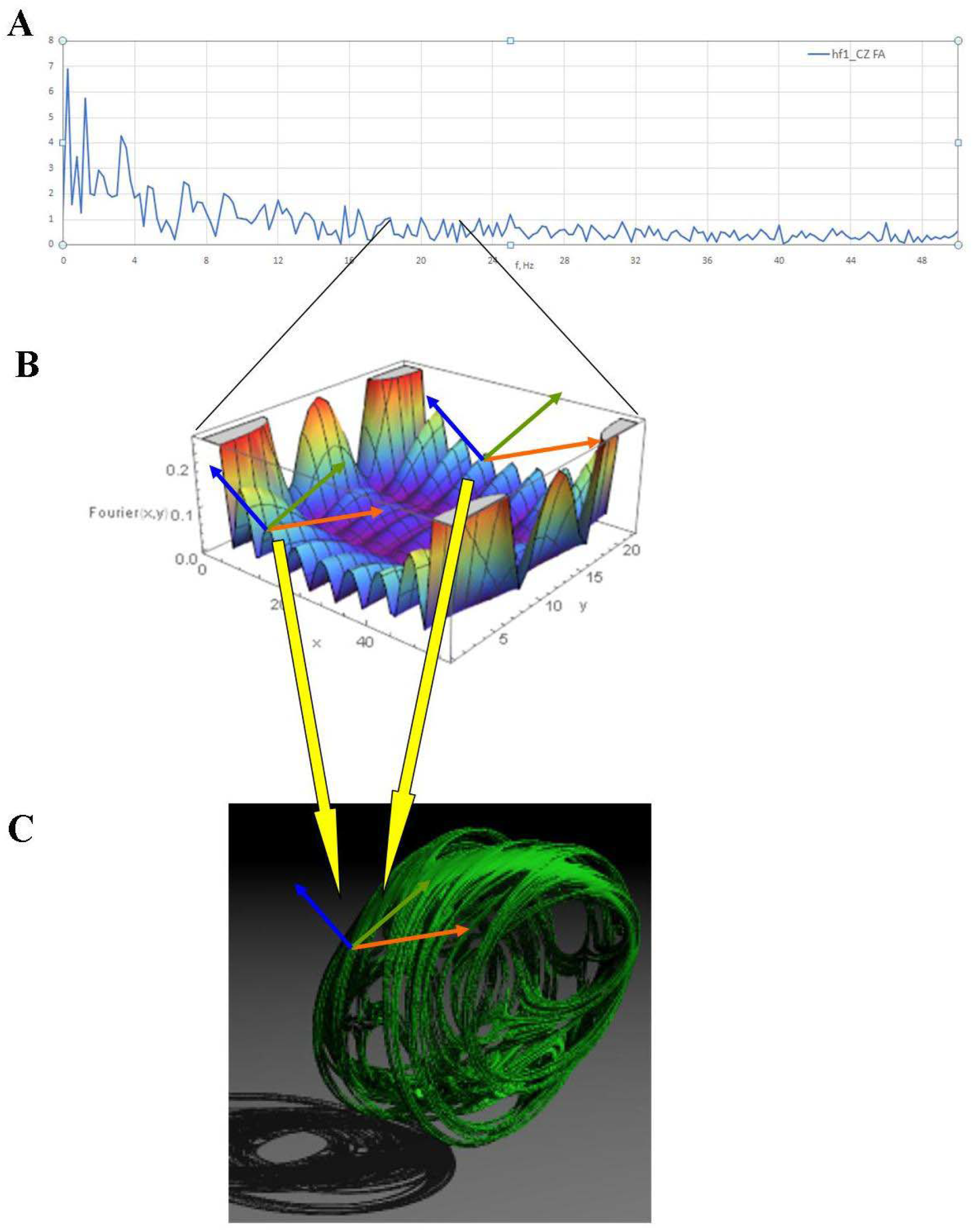
Quaternion simulation of fractal, non-uniform orthographic projection of brain signals onto a fragmented sphere. **Figure 3A**: Two-dimensional Fourier analysis of an EEG trace. The x axis displays the frequencies in Hertz, while the y axis the amplitude in mV. The plot is equipped with scale-free dynamics. **Figure 3B**: three-dimensional magnification of a short trace of Figure 3A. **Figure 3C**: view of the orthographic projection of a brain signals patch on continuous curves represented by green threads and their shadows via a quaternion. Note that two triads with matching description in three dimensions (i.e., equipped with the same orientation of the three axes) project to a single triad in four dimensions. The interdimensional mapping is illustrated by the two yellow arrows.

To achieve the three-dimensional fractal quaternionic orthographic projections of brain signal patches treated with Fourier analysis,we performed simulations depicting the temporal progression of there scale-free dynamics. Indeed, as stated above, an EEG Fourier analysis is characterized by power laws behaviour. **Figure 4** portrays simulations of fourdimensional quaternionic orthographic projections of fractal paths in the brain. In particular, the bumps illustrated inthe four-dimensional fractal quaternionic plot of **Figure 4E** represent varying brain signal amplitudes moving along surfaces inside the brain.

**Figures4.**
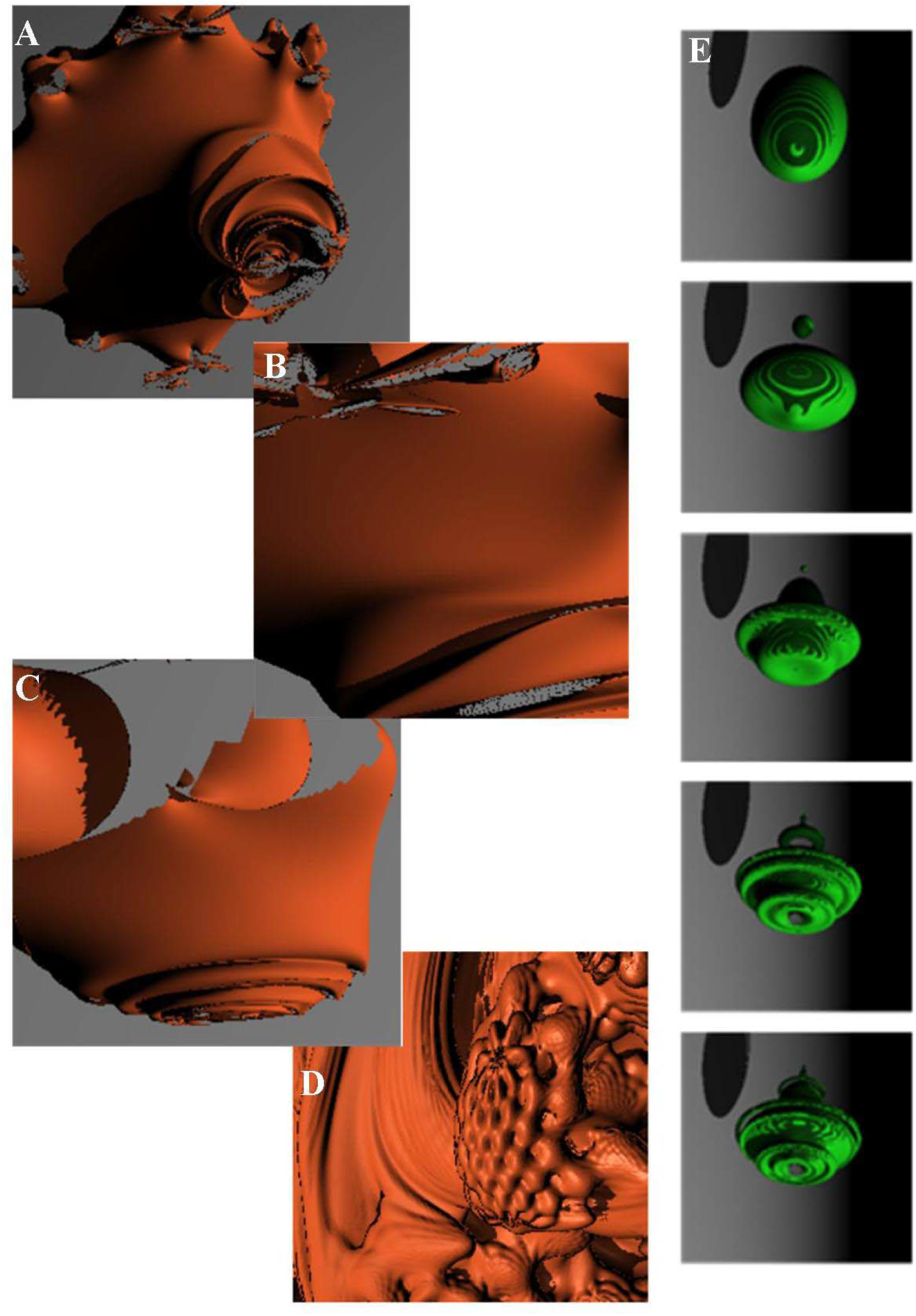
**A-D**. Simulations of orthographic projections of brain signal patches onto the surface of a sphere via quaternions. Every picture stands for a quaternion surface closeup, where magnifications of the fractal structure of the hypersphere, depicted as three-dimensional views of the quaternion, can be seen. To make an example, **Figure 4A** illustrates a planetary view of a non-uniform orthographic projection of brain signals onto a fragmented sphere. **Figure 4E**: fractal-view (from the top to the bottom:) of the orthographic projection of temporal sequences of brain signals patches on continuous curves represented by pulsating (vibrating) green threads and their shadows via a quaternion. This sequence stands for a spacetime view of the quaternionic mappings and encompasses a hidden time scale for the sequence of orange orthographic projections.

## CONCLUSIONS

We simulated quaternion orthographic projections of brain signals, using fractal traces from Fourier analysis of EEG plots. The recent, exciting insights in quaternion-valued neural networks, such as, e.g., the ones suggested by Li and Wang (2018) and by Oliveros-Muñoz et al. (2013), pave the way to test an intriguing hypothesis: the recently-proposed claimthat the brain activity takes place in multidimensional spaces. It is noteworthy that, apart from the quaternionic maps described in this paper, another method is available to assess higher dimensional activities of nervous networks. Indeed, it has been recently suggested (Tozzi 2019) that hidden spatial dimensions of nervous activity could be tackled in terms of oscillations inside a two-dimensional superlattice equipped with quantum Hall effects. In this peculiar apparatus, the superimposition of waves of different wavelengths gives rise to both (two-dimensional) linear and (four-dimensional) nonlinear dynamics (Lohse et al., 2018; Zilberberg et al., 2018). Both the approaches, i.e., quantum Hall effects and quaternion-valued neural networks, would allow to transfer mathematical/physical issues to the realm of neuroscience, in order to: a) describe the real multidimensional brain dynamics and b) operationalize the feasibility of a synthetic nervous network equipped with four spatial dimensions (plus time), instead of the classical three (plus time). Importantly, quaternionic views of brain activity lead to a descriptive proximity (Di Concilio et al., 2018) characterization of the activities of different brains that are descriptively close. This is made possible by the recent introduction of approximate descriptive proximity (Peters, 2020), which facilitates computation of the distance between feature vectors used to represent and describe the shapes of brain signal patches. Thanks to the role of quaternions introduced in this paper, quaternion-based feature vectors provide a formal basis for the approximate descriptively proximal assessment of the closeness of spatially separated brain activities either by the same brain or by different brains at different times.

A crucial question is still unsolved: apart from the assessment of the multidimensional brain model, may quaternionic maps provide other insights that could be useful to describe the conventional three-dimensional neural dynamics? The answer is positive: indeed, quaternionic mapping stands for a versatile technique that has been widely used in different experimental contexts. To make a few examples, Hanson and Thakur (2012) used quaternion to expose global spatial relationships among amino acid residue structures within proteins: this allowed them to assess global residue alignment in crystallographic data and statistical orientation properties in Nuclear Magnetic Resonance. The visualizations resulting from quaternion spaces enables comparisons of absolute as well as relative orientations, allowing, e.g., to ascertain the stereochemical nature of proteins, or to visualize global and local residue orientation properties of nucleic acids.

As stated before, a quaternion mapping embodies the full three degree-of-freedom transformation from the threedimensional identity frame triad to a four-dimensional frame triad. This means that a quaternionic representation is much simpler than the usual representation using a triple of orthogonal three-dimensional vectors (Hanson and Thakur, 2012). The three spatial coordinates of every point inside a three-dimensional plot become a single point inside a four-dimensional hypersphere. In particular, the points located close to each other on the hypersphere’s surface display similar coordinates in three dimensions: in other words, these points are equipped with matching description. This allows quaternions to naturally expose, e.g., global similarities among spatially far apart residues in a protein complex, no matter how near or how distant, extending the analysis to far between components of multi-part structures (Hanson and Thakur, 2012). Quaternion mappings onto a hypersphere disclose properties of the dynamic systems’ spatial orientation, making easier to detects hidden features such as global or broken symmetries, zones of speed convergence, domains of attraction inside three-dimensional projections, convergence among apparently random paths (Oliveros-Muñoz et al., 2013). Therefore, quaternions provide an alternative methodology to the widely diffused methods of time-lagged mutual information and autocorrelation functions that have been used to evaluate the periodic features of the continuous EEG signal (von Wegner 2018; von Wegner et al., 2018). The last, but not the least, an intriguing hypothesis can be raised: as well as Fruchart et al. (2020) investigated hidden non-commutative dualities (i.e., mathematical mappings that reveal links between apparently unrelated systems) in twisted kagome lattices, quaternionic movements could detect neural responses not predicted by standard symmetry analysis, in order to assess self-dualities in brain dynamics that are dictated by non-commuting responses.

In sum, mapping EEG oscillations to an S^3^ hypersphereis accomplished with orthographic projections of vectors inherent in brain signal patches via quaternions: this relatively straightforward approach leads to the possibility to enlighten hidden symmetries, in particular in nervous neurodata.

